# Macrophage to Myocyte Mitochondrial Transfer in the Myometrium: A Novel Mechanism for the Initiation of Labor

**DOI:** 10.64898/2026.08.17.745259

**Authors:** Lubna Nadeem, Amit Sharma, Sharanya Shankar, Eduardo Aguiar-Cabeza, Laurence Pelletier, Andrea Juriscova, Oksana Shynlova, Stephen Lye

## Abstract

Labor initiation involves influx of peripheral monocytes into uterine tissue, where they differentiate into macrophages (Macs) and polarize toward pro-inflammatory (M1) or anti-inflammatory (M2) phenotypes. Crosstalk between uterine myocytes (MYO) and Macs is implicated in myometrial activation, but the underlying mechanisms remain unclear. Using immunofluorescence and live-cell imaging, we show that M1-Macs (but not M2-Macs) form tunneling nanotubes (TNT) with MYOs, enabling direct, unidirectional mitochondrial transfer. TNT-mediated mitochondrial transfer enhances ATP production and 20α-HSD expression in MYO. Functionally, M1-Macs induce intracellular and functional progesterone (P4) withdrawal in MYOs via 20α-HSD mediated P4 metabolism and increased PR-A phosphorylation. In vivo, transfer of monocytes, carrying dendra-labelled mitochondria in pregnant mice confirm their myometrial recruitment, differentiation, polarization to M1-Macs, and M1-Mac/MYO mitochondrial transfer. Increased M1-Macs and mitochondrial transfer to MYOs are observed before labor onset, implicating M1-Macs/MYO interaction as key regulator of labor initiation and potentially a therapeutic target to prevent preterm birth.

## Introduction

The mechanisms governing the initiation of labor are complex and involve coordinated interactions between the maternal reproductive and immune systems. Our previous studies have demonstrated a strong correlation between maternal immune cell infiltration into the uterine smooth muscle (myometrium) and the initiation of labor, both at term (TL) and preterm (PTL) ^1, 2, 3, 4^. We established that the myometrium functions as an immune-regulatory tissue, capable of producing cytokines and chemokines that activate maternal innate immunity and recruit peripheral immune cells, particularly monocytes and neutrophils, facilitating their trans-endothelial migration. Infiltrating monocytes differentiate into macrophage (Macs) within the myometrium and exhibit functional plasticity by shifting between pro-inflammatory M1 and anti-inflammatory M2 phenotypes depending on the local microenvironment.

Accumulated evidence suggests that withdrawal of progesterone (P4) is essential for labor onset. P4 maintains the myometrium in a quiescent, non-contractile state by suppressing the transcription of contraction-associated proteins (CAPs), which are essential for initiating myometrial contractions and labor. In most species, P4 levels decline at term and hence trigger labor induction. However, P4 levels remain elevated in the systemic circulation throughout human pregnancy. We resolved this paradox by showing that P4 levels are locally/intracellularly reduced within uterine MYOs during labor, due to catabolism by 20alpha-hydroxysteroid Dehydrogenase (20α-HSD), leading to unliganding of P4 receptors (PR-A and PR-B), and dominance of PR-A. Crucially, in its unliganded state, PR-A switches from a repressor to an activator of CAP gene transcription ^5^. We have recently reported that inflammatory stimuli can induce intracellular P4-withdrawal in MYOs ^6^ and that M1-Macs contribute to this process ^7^.

Emerging evidence suggests a potential role of Macs in orchestrating the maintenance of pregnancy and even the timing of labor onset. Our studies in rodents show that Macs are present within the uterine tissues throughout pregnancy, increasing in late gestation and during both TL and PTL ^3, 8^, Macrophage depletion in rodent models prevents infection-induced preterm birth ^9^ and cervical remodeling ^10^, while the adoptive transfer of M2-Macs protects against PTL and improves neonatal outcomes in a model of sterile intra-amniotic inflammation induced by HMGB1 ^11^. Consistent with these findings, human and mouse studies show a predominance of M2-Macs during late gestation and their decline at TL, suggesting a role in maintaining uterine quiescence ^12^. In contrast, M1-Macs are increased in the decidua and the myometrium in association with labor onset^13, 14^. Recent evidence suggests crosstalk between activated Macs and uterine MYOs resulting in increased reactive oxygen species (ROS) ^15^ however, the mechanism underlying this crosstalk is unknown. Understanding the mechanisms that govern Mac-MYO communication during pregnancy is essential for successful development of therapeutic strategies to prevent PTB.

In this study we hypothesize that the pro-inflammatory M1-Macs directly interact with and activate uterine MYOs inducing 20α-HSD mediated intracellular P4-withdrawal, dominance of PR-A and CAP gene expression. We tested this hypothesis using human Mac-MYO co-culture and examined the Mac-MYO interaction in-vitro through time-lapse imaging and in-vivo by investigating monocyte infiltration, differentiation and Mac polarization in pregnant murine myometrium from mid-gestation to post-partum. Our study provides novel insights into labor initiation and implicates mitochondrial transferred to uterine myocytes from M1-Macs as one of the mediators in this process.

## Materials and Methods

### Ethics

All human tissue and blood sample collections were approved by the Sinai Health Research Ethics Board (SH REB #04-0024E and #12-0007-E). Informed consent was obtained from women donors of myometrial tissues involved in this study prior to sample collection. Myometrial biopsies were collected from healthy pregnant women at term with no signs of labor (TNL) undergoing elective caesarean section for breech presentation or repeat C-section, or laboring (TL) undergoing an emergency caesarian section due to fetal distress. Peripheral blood was collected from healthy pregnant women in their third trimester (37–40 weeks of gestation) into EDTA vacutainer tubes.

All mice-related experiments and procedures were approved by the Animal Care Committee of The Centre for Phenogenomics (TCP, Toronto, Canada) (AUP #0164H) and conducted in accordance with the guidelines of the Canadian Council on Animal Care. Mice were housed in a pathogen-free, humidity-controlled facility under a 12 h light/12 h dark cycle, with ad libitum access to food and water.

### Myocyte Cell lines and cell culture

***a) Primary human myometrial cell lines*** were derived from the myometrial biopsies collected from non-laboring term pregnant women (TNL) undergoing elective caesarian section for breech presentation or repeat section, or laboring women (TL) undergoing emergency caesarian section due to fetal distress. Myometrial tissues were collected in ice-cold HBSS containing Ca²⁺ and Mg²⁺, supplemented with 2.5% HEPES (Wisent, Montreal, QC, Canada) and 1% penicillin/streptomycin (Pen/Strep, Gibco^TM^, Thermo Fisher Scientific, Waltham, MA, USA). Cell isolation was performed as previously described ^16^. Briefly tissues were cleared of blood vessels, minced (∼1 mm³), and washed sequentially in HBSS+/+ and HBSS−/−. Samples were digested for 1 h at 37 °C in a rocking water bath using an enzymatic solution containing 10% fetal bovine serum (Wisent, Montreal, QC, Canada),1 mg/mL collagenase II, 1 mg/mL bovine serum albumin, 0.15 mg/mL DNase I, and 0.1 mg/mL trypsin inhibitor (Sigma-Aldrich, Oakville, ON, Canada). Digested tissue was mechanically dissociated in ice-cold dissociation solution (HBSS−/−, with 10% FBS (Wisent, Montreal, QC, Canada), 1 mg/mL BSA (Wisent, Montreal, QC, Canada), filtered through 70-micron filter, and residual tissue was subjected to a second digestion. Combined cell suspensions were centrifuged, washed in DMEM with 10% fetal bovine serum, passed through a 23 G x 3/4″ needle, and resuspended in complete growth medium (DMEM medium supplemented with 20% fetal bovine serum and 1% Pen/Strep). Cells were seeded in 10 cm culture dishes and maintained at 37 °C in a 20% oxygen incubator.
***b) Immortalized human myometrial cell line***; hTERT-HM^A/B^ (derived from hTERT-HM parent cell line ^17^, with inducible expression of progesterone receptors (PRs: PR-A and PR-B) was a gift from Dr Sam Mesiano ^18^. The cell line was maintained in DMEM/F12 (Gibco, Sigma Aldrich, USA) supplemented with 10 % fetal bovine serum (Wisent, Montreal, QC, Canada), 1% Pen/Strep (Lonza, Basel, BS, Switzerland), and the selection antibiotics (0.1 mg/mL geneticin, 0.001 mg/mL hygromycin, and 0.005 mg/mL blasticidin from Life Technologies).
***c) Primary mouse myometrial cell lines*** were derived from pregnant C57-BL/6 or Mitodendra2-C57-BL/6 mice ^19^ at mid-gestation (GD15) using the same protocol as described above for human myometrium. Cells were similarly cultured in DMEM medium supplemented with 20 % FBS and 1 % Pen/Strep.

### Source of Macrophages

#### a) Peripheral blood derived monocyte isolation from term pregnant women

Peripheral blood samples were collected from third-trimester pregnant women (37–40 weeks of gestation). Monocytes were isolated using the RosetteSep Human Total Monocyte Enrichment cocktail (STEMCELL Technologies, Vancouver, BC, Canada) according to the manufacturer’s instructions. Briefly, RosetteSep cocktail supplemented with EDTA (50 µL/mL blood) was added to whole blood and incubated at room temperature for 20 min. Samples were diluted 1:1 with PBS containing 2% FBS and layered over Ficoll (Sigma-Aldrich, Oakville, ON, Canada), followed by centrifugation at 1200 rcf for 20 min. The monocyte-enriched layer was collected, washed twice with PBS, and resuspended in RPMI supplemented with 10% FBS (Wisent Inc., Montreal, QC, Canada).

#### b) THP-1 cell line

The human monocytic leukemia cell line THP-1 (ATCC® TIB-202™) was obtained from a certified cell repository and maintained according to recommended guidelines. Cells were cultured in RPMI-1640 medium (Gibco^TM^, Thermo Fisher Scientific, Waltham, MA, USA) supplemented with 10% FBS and 1% penicillin-streptomycin.

#### c) Murine peripheral blood isolation of Monocytes

Peripheral blood samples were collected from pregnant C57-BL/6 or Mitodendra2-C57-BL/6 mice ^20^ at mid-gestation (GD15). Monocytes were isolated using the EasySep^TM^ Mouse Monocyte Isolation Kit (STEMCELL Technologies, Vancouver, BC, Canada) according to the manufacturer’s instructions. Briefly, red blood cell lysis was conducted on ice for 15 minutes using erythrocyte lysis buffer (QIAGEN, Hilden, Germany), followed by centrifugation at 300 g for 6 minutes. Cell pellet was resuspended in 500 µl of Ca^++^/Mg^++^ free PBS supplemented with 2 %FBS, 1 mM EDTA). Negative selection was conducted using the company provided cocktail and magnetic beads. The isolated cells were cultured in RPMI supplemented with 10% FBS and 1% penicillin-streptomycin.

### Differentiation of Monocytes to Macrophages

Primary monocyte differentiation was carried out using the method described by Karina et al. ^21^ with slight modifications. Monocytes were allowed to differentiate into macrophages for 10 days in media containing granulocyte-macrophage colony-stimulating factor (GM-CSF, Sigma-Aldrich, St. Louis, MO, USA) for M1- and Macrophage colony-stimulating factor (M-CSF) for M2-Macs, at a concentration of 100 ng/mL. The media was refreshed every three days. THP-1 monocytes were differentiated into Macs according to the protocol outlined by Shiratori et al. ^22^ with some modifications. Specifically, the THP-1 monocytes were treated with 50 ng/mL of Phorbol 12-myristate 13-acetate (PMA) for 48 h followed by incubation in complete medium for an additional 72 h.

### Polarization of Macrophages

Primary human Macs (hMacs) and murine Macs (mMacs) were polarized into M1-Macs with LPS (100 ng/mL), and IFNɣ (25 ng/mL), and into M2-Macs with IL-13 (25 ng/mL,) and IL-4 (25 ng/mL) treatments for 48 h. Lipopolysaccharide (LPS) was purchased from Sigma-Aldrich, and all other reagents were obtained from R&D Systems Inc., Minneapolis, MN, USA. Macs were harvested using Macrophage Detachment Solution Promocell (Sigma Aldrich), washed with RPMI supplemented with 10% FBS (Wisent Inc., Montreal, QC, Canada), and pelleted by centrifugation at 200× g for 5 min.

THP-1-derived Macs were polarized to hM1-Macs by incubating with 50 ng/mL of LPS and 20 ng/mL of IFN-γ for 48 h. For M2 polarization, THP-1-derived Macs were treated with 20 ng/mL of IL-4 for 48 h. Polarization was verified by real-time PCR using specific markers for M1 (*CCL2*, *IL6*, *IL1B*, and *TNFA*) and M2 (*CD206* and *PPARG*) Macs, with expression levels normalized to three housekeeping genes (*HPRT*, *SDHA*, *TBP*) (**Supplemental Figure 1**). Primer sequences are provided in **Supplemental Table S1.**

### Time Lapse imaging

Primary human (hMYO) or mouse (mMYO) myometrial cells were seeded in Nunc™ Lab-Tek™ II chambered cover glass (Thermo Fisher Scientific, Waltham, MA, USA). Cells were stained with either the plasma membrane marker wheat germ agglutinin (WGA; Thermo Fisher Scientific) or the F-actin probe SPY650-FastAct™ (Spirochrome, Schaffhausen, Switzerland). M1-Macs were stained with MitoTracker™ CM-H2XRos dye alone or in combination with the plasma membrane dye Vybrant™ DiD (Thermo Fisher Scientific). Labelled M1-Macs were co-cultured with MYOs in the presence of 100 nM progesterone (Sigma-Aldrich, St. Louis, MO, USA). Specific staining conditions for each experiment are detailed in the corresponding figure legends. Time-lapse imaging was performed using a Nikon A1R laser-scanning confocal microscope equipped with a 40× water-immersion objective (Apo LWD 40× WI λS DIC N2). Images were acquired over extended imaging periods (14-20 hours), with acquisition intervals between 6-10 minutes. Confocal images were denoised using Nikon NIS-Elements software, and movies were generated as maximum-intensity projections.

### Protein extraction and Immunoblotting

Protein extracts were prepared as previously described ^6^. Briefly, cells were washed with ice-cold phosphate-buffered saline (PBS) and lysed on ice in buffer containing Tris-HCl (1 M, pH 6.8), 10% sodium dodecyl sulfate (SDS), 5% glycerol, and 1% Halt protease and phosphatase inhibitor cocktail (Thermo Fisher Scientific, Waltham, MA, USA). Lysates were transferred to pre-chilled tubes, vortexed for 15 s, and incubated on ice for 10 min. Samples were then sonicated (XL-2000 Sonicator, Misonix Inc., Farmingdale, NY, USA) at 2.5 dB for 10 s, followed by incubation on ice for 5 min. Lysates were subsequently heated at 95 °C for 5 min, cooled on ice for an additional 5 min, and centrifuged at 14,000 rpm for 25 min at 4 °C. The clarified supernatants were collected into pre-chilled 1.5 mL microcentrifuge tubes and stored at -20 °C until further use. Protein concentrations were determined using the Pierce BCA Protein Assay Kit (Thermo Fisher Scientific, Waltham, MA, USA) according to the manufacturer’s protocol.

Equal amount (µg) of proteins was resolved on 10–12% TG-SDS polyacrylamide gels under reducing conditions. BLUelf Prestained Protein Ladder (GeneDireX, Taoyuan, Taiwan) was loaded as a molecular weight reference. Proteins were transferred onto polyvinylidene difluoride (PVDF) membranes using a Trans-Blot system (Bio-Rad, Mississauga, ON, Canada). Membranes were blocked for 1 h at room temperature in 5% non-fat milk (blocking buffer) prepared in Tris-buffered saline with 0.1% Tween-20 (TBS-T), followed by overnight incubation at 4 °C with primary antibodies diluted in 1% Bovine Serum Albumin (BSA) prepared in TBS-T or milk blocking buffer. The following day, membranes were washed three times with TBS-T (10 min each) and incubated with horseradish peroxidase (HRP)-conjugated secondary antibodies (Thermo Fisher Scientific, Waltham, MA, USA; 1:5000 dilution) for 1 h at room temperature. After three additional washes with TBS-T, immunoreactive bands were detected using chemiluminescence reagents (Thermo Fisher Scientific, Waltham, MA, USA) and imaged with ChemiDoc (BioRad, Hercules, California, USA). Details of primary antibodies used in this study are provided in **Supplementary Table S2**. Band intensities were quantified using Image Lab software (version 4.0.1 build 6; Bio-Rad), and protein expression levels were normalized to housekeeping proteins tubulin or ERK2 or total protein when run on a stain-free gel.

### Progesterone Quantification by ELISA

hTERT-HM^A/B^ cells (MYOs) were seeded in 96-well plates at the density of 20,000 cells per well. THP-1–derived M1 or M2 macrophages were co-cultured with MYOs at a ratio of 1:10 (Macs:MYOs) in serum-free medium supplemented with 1% ITS and treated with 10 nM P4. MYO monocultures served as controls, and a positive control was included by treating MYOs with lipopolysaccharide (LPS, 1 µg/mL), which has previously been shown to reduce P4 levels in vitro ^6^. Conditioned medium was collected 24 h after treatment. P4 levels in the conditioned medium were measured using a progesterone ELISA kit, Catalogue # 582601 (Cayman Chemical, Ann Arbor, Michigan, USA) according to the manufacturer’s instructions. Briefly, standards and samples were prepared in ELISA buffer and added in duplicate to wells of a 96-well plate pre-coated with goat anti-mouse IgG. Progesterone-acetylcholinesterase (AChE) tracer and progesterone-specific monoclonal antibody were added to each well, followed by incubation for 2 h at room temperature on an orbital shaker to allow competitive binding. Wells were washed to remove unbound reagents, and Ellman’s reagent was added for color development. Plates were incubated in the dark at room temperature for 60–90 min, and absorbance was measured at 420 nm using a microplate reader (Tecan Infinite 200 Pro, Männedorf, Switzerland). P4 concentrations were calculated from a standard curve using the manufacturer-provided analysis spreadsheet at: “https://www.caymanchem.com/analysisTools/elisa”.

### ATP Quantification Assay

Cellular ATP levels were measured using the ATPlite Luminescence Assay System (PerkinElmer, Catalogue # 6016736) according to the manufacturer’s instructions. Briefly, MYOs were seeded in white opaque 96-well plates at the indicated density and allowed to adhere overnight. The following day, MYOs were co-cultured with THP-1–derived M1 or M2-Macs at a ratio of 1:10 (Macs:MYOs). MYOs, M1 and M2-Macs were also maintained as monocultures at the same seeding density for comparison. After 24 h of co-culture, an equal volume of cell lysis solution was added to each well, followed by incubation for 5 min with orbital shaking to ensure complete cell lysis. Subsequently, substrate solution containing luciferase and luciferin was added, and plates were incubated in the dark for 10 min to allow stabilization of the luminescent signal. Luminescence, proportional to intracellular ATP levels, was measured using a microplate reader (Tecan Infinite 200 Pro, Männedorf, Switzerland). Background signal from media-only wells was subtracted, and ATP levels were quantified relative to a standard curve.

### Mitochondrial Isolation and treatment

Mitochondria were isolated from THP-1 or mitoDendra-Monocytes-derived M1-Macs using the Mitochondria Isolation Kit for Cultured Cells Catalogue # 89874 (Thermo Fisher Scientific, Waltham, MA, USA) according to the manufacturer’s instructions. Briefly, cells were trypsinized, washed with ice-cold PBS, and pelleted by centrifugation at 1400 rpm for 5 min. PBS was removed, and cell pellets were resuspended in Reagent A supplemented with protease inhibitors and incubated on ice for 2 min. Reagent B was then added, and cells were gently vortexed for 5 s, followed by further incubation on ice for 5 min to facilitate membrane disruption. Reagent C was subsequently added, and samples were centrifuged at 700 × g for 10 min at 4 °C to pellet nuclei and intact cells. The resulting supernatant was transferred to a fresh tube and centrifuged at 3,000 × g for 15 min at 4 °C to pellet mitochondria. The cytosolic fraction was collected, and the mitochondrial pellet was washed once with Reagent C and centrifuged at 12,000 × g for 5 min. The supernatant was discarded, and the mitochondrial pellet was resuspended in culture medium (without FBS) and used immediately. The purity of the isolation procedure was assessed by Western blotting, in which total cell lysates from mitoDendra-M1-Macs were compared with the cytosolic fraction depleted of mitochondria (C-Mito) and the isolated mitochondrial fraction (Mito). An anti-Dendra antibody was used to verify the presence of mitoDendra-derived mitochondria (**Supplemental Figure 2**).

Isolated mitochondria were transferred to MYO monolayer through mitoception procedure as described earlier ^23^. Briefly, mitochondria were added to monolayers of hTERT-HM^A/B^ cells (hMYOs) cultured in serum-free medium supplemented with 1% ITS and treated with 100 nM progesterone (P4). The culture plates were then centrifuged at 1,500 g for 15 min at 4°C followed by placing in 37°C cell incubator for 24 h. Cells were washed with cold PBS thrice and harvested for protein analysis by Western blotting. To validate the uptake of mitochondria by MYOs, mitochondria were similarly isolated from mitoDendra-M1-Macs expressing Dendra (green fluorescence) and added to mMYOs pre-stained with SPY650-FastAct™ (red). Cells were washed and imaged 24 h post-mitochondrial transplant to assess mitochondrial uptake by mMYOs. Representative data are shown in **Supplemental Figure 2.**

### Adoptive transfer of *mitoDendra*-Monocytes in pregnant mice

Peripheral blood monocytes (mitoDendra-monocytes carrying Dendra-labelled mitochondria) were isolated from pregnant MitoDendra2-C57BL/6 mice on gestational day (GD) 15. Monocytes were resuspended in sterile saline, and 4 × 10⁴ to 1 × 10⁵ cells per dam were administered to pregnant wild-type (WT) C57BL/6 mice via tail vein injection on GD15. Uteri from injected WT mice were collected at 24-h intervals throughout gestation; i.e. GD16, GD17, GD18, and GD19/term not in labor (TNL), GD19/term labor (TL) and GD19/postpartum (PP). Tissues were cryopreserved in optimal cutting temperature (OCT) compound.

### Multiplex-Immunostaining of murine tissues

Serial cryosections of the uterus from WT mice that received mitoDendra-monocytes were initially subjected to detection of Dendra using a proximity ligation assay (PLA). PLA amplification (bright red), performed using an anti-Dendra antibody in conjunction with complementary plus and minus probes, enabled specific detection and signal amplification, ensuring that Dendra signals were distinct and above the threshold of background autofluorescence. The same tissues were subsequently immunostained with the pan-macrophage marker F4/80 (Cy5; bright pink) in combination with either the M1 marker CD38 or the M2 marker EGR2 (Alexa Fluor 488; green), which are established markers of murine M1 and M2 macrophage phenotypes, respectively as reported earlier ^24^. Nuclei were counterstained with 4′,6-diamidino-2-phenylindole (DAPI; blue) (**Supplemental Figure 3).** In corresponding serial sections, co-staining for smooth muscle actin (SMA), F4/80, and Dendra was also performed. Parallel IgG controls were prepared for each antibody combination. Details of all antibodies used are provided in **Supplementary Table S2.**

### Statistical analysis

All data were assessed for normality using the Shapiro-Wilk test. For comparisons between two groups, unpaired t-tests were applied to normally distributed data. Outlier detection was conducted using the ROUT or Grubbs’ test, assuming a normally distributed population. For data comprising more than two groups with a normal distribution, statistical significance was evaluated using one-way ANOVA followed by Dunnett’s post hoc multiple comparisons test, and for data involving multiple groups with two independent variables, two-way ANOVA with Dunnett’s multiple comparisons post-test was employed. All statistical analyses were performed using GraphPad Prism (Version 10/11; GraphPad Software Inc., CA, USA). Statistical significance was denoted as p < 0.033 (*), p < 0.002 (**), and p < 0.001 (***).

## Results

### M1-polarized Macrophages physically connect with Myocytes through tunneling nanotubes

We examined cell-cell connectivity between human myometrial smooth muscle cells (hMYOs) and Macs using a “parachute assay”. Human M1-polarized Macs (hM1-Macs), derived from peripheral blood monocytes of term pregnant women or from monocytic cell line THP-1 were loaded with calcein dye and co-cultured with primary hMYOs derived from myometrial biopsies from TNL women or with myometrial cell line hTERT-HM^A/B^ at a ratio of 1:100. Fluorescent microscopy revealed physical connections between hM1-Macs and hMYOs within two hours of co-culture. Distinct tubular membrane projections were seen extending from the hM1-Macs toward the surrounding hMYOs (**Figure 1A**) along with the transfer of calcein dye from the hM1-Macs to the hMYOs, indicating the presence of functional open channels facilitating intercellular dye transfer (**Figure 1A, Supplemental Figure 4**). These projections were classified as tunneling nanotubes (TNTs), based on the previously established criteria ^25, 26^. Specifically, TNTs are defined as plasma membrane projections that physically connect cells either in close proximity or at relatively far distance. Key characteristics of TNT include: 1) substrate independence; 2) positivity for F-actin; 3) variability in length and thickness; 4) the capacity for transcellular transfer of cellular cargo. The protrusions observed between hM1-Macs and hMYOs fully conform to these criteria. The substrate independence of TNT projections originating from hM1-Mac (marked with asterisk in **Figure 1A**), as well as transfer of calcein dye to hMYO are shown in **Figure 1A**. The hM1-Macs established TNT-mediated connections with nearby as well as distant hMYOs (**Figure 1B, C, Supplemental Video 1, 2**). These TNTs show positive staining for F-actin and have variable lengths and diameters (**Figure 1C, D**). We also observed the bulge-like structures referred to as “gondolas”^27, 28^, along the nanotubes connecting the two cells suggesting active transport of cellular cargo between them **Figure 1C** (inset image, yellow arrow).

**Figure 1:**
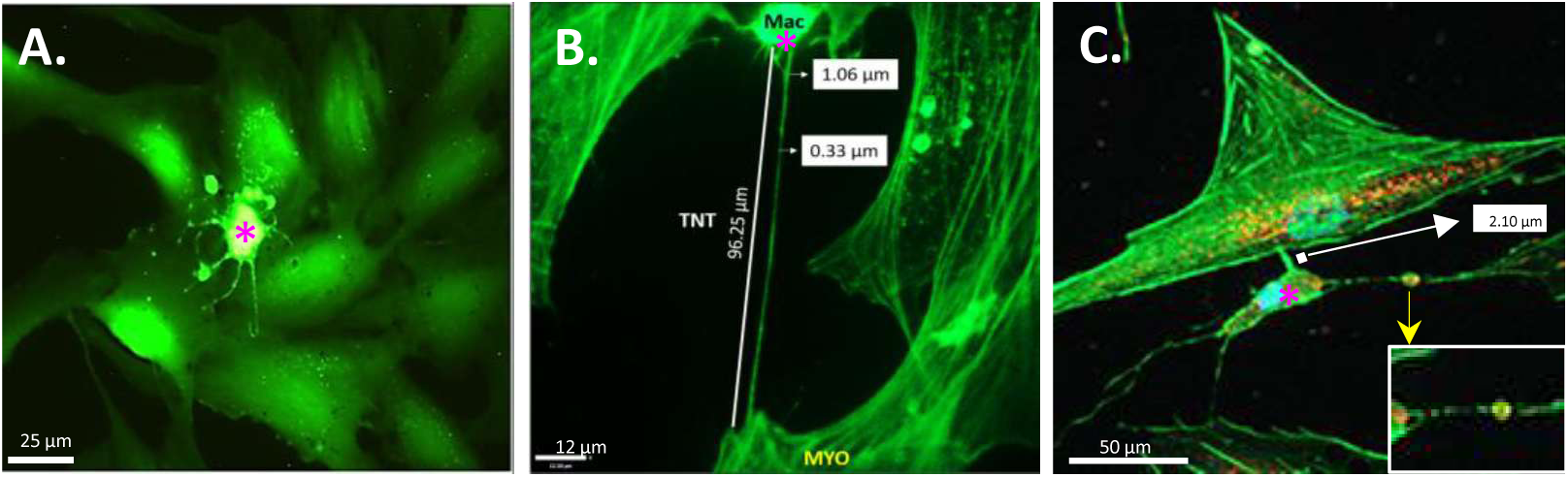
M1 marophages physically connect with myocytes via tunneling nanotubes. **A)** THP-1-derived human M1-macrophages (hM1-Macs) were labelled with calcein-AM and co-cultured for 2 hours with unlabelled hTERT-HM^A/B^ myocytes (hMYO). hM1-Macs (indicated by asterisk) exhibited membrane protrusions that extended towards neighboring myocytes. Transfer of green calcein fluorescence into myocytes indicates functional M1-Mac/MYO connectivity. **B–C) Membrane protrusions exhibit features consistent with tunneling nanotubes**. PBMC-derived hM1-Macs (asterisks) were co-cultured with primary hMYO for four hours and stained with phalloidin-Alexa Fluor 488 (F-actin marker, green). The protrusions show F-actin positivity, substrate independence, variable thickness and length, and evidence of cargo transfer. In panel **(C)** hM1-Macs (asterisks) were preloaded with MitoTracker dye (orange-red), and mitochondrial transfer to MYO cells was detected via tunneling nanotubes. A mitochondrion-containing “gondola” is shown in the inset (yellow arrow).

To further examine this intercellular cargo transfer, we used MitoTracker red dye to label the hM1-Macs. The mitochondria-labelled hM1-Macs were then co-cultured with hMYOs. After four hours of co-culture, cells were fixed and stained with phalloidin conjugated with Alexa Fluor-488 to visualize TNTs by green F-Actin. We observed the formation of TNT bridges between hM1-Macs (marked with pink asterisk, **Figure 2A**) and hMYO (**Figure 2C**) and detected labelled mitochondria within the TNT bridge (**Figure 2B**, white arrows) and in hMYO (**Figure 2A, B**), indicative of open-ended TNTs connection between cells.

**Figure 2:**
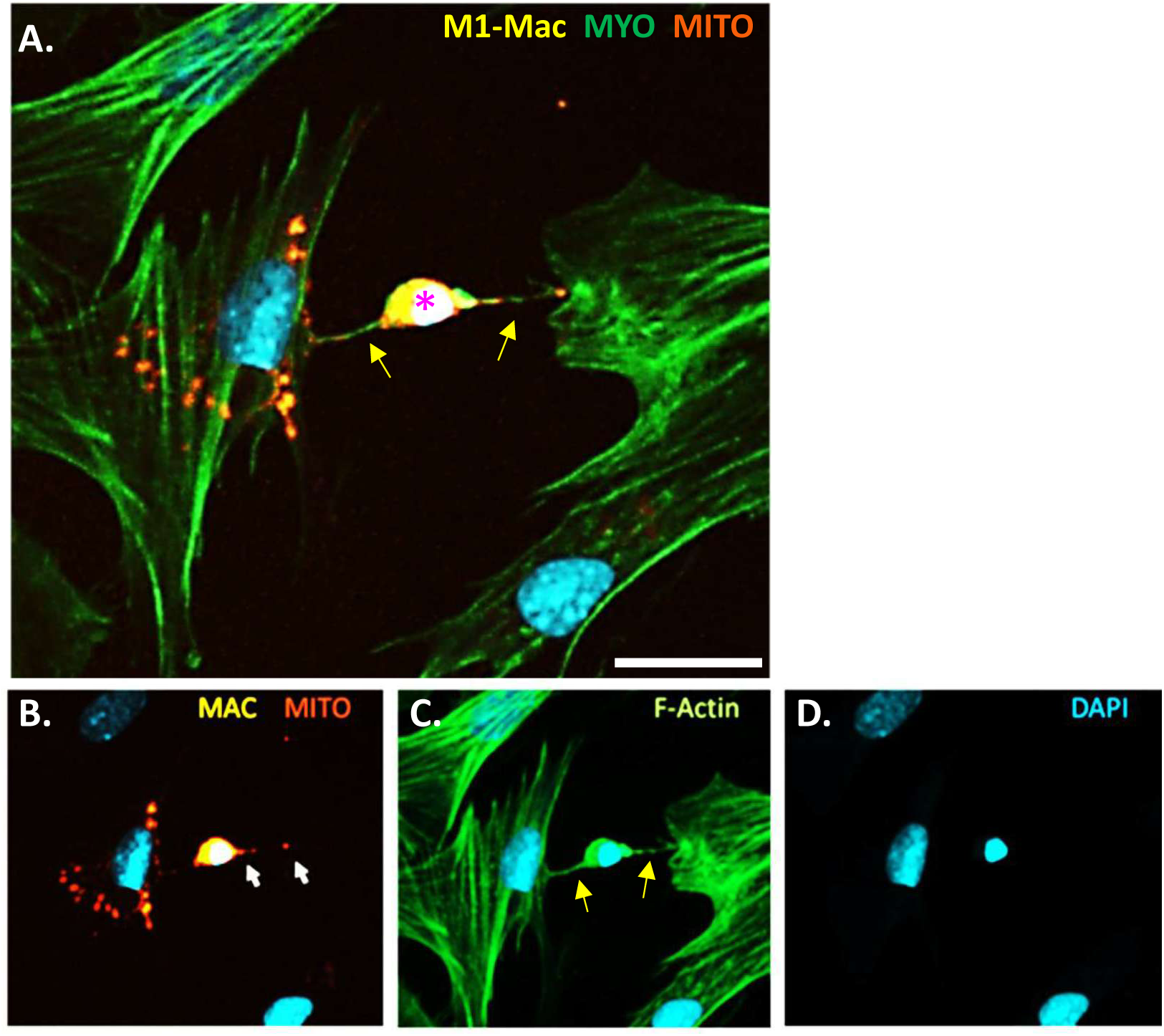
M1 macrophages transfer mitochondria to the myocytes through tunneling nanotubes. Human PBMC-derived M1 macrophages (hM1-Mac, indicated with asterisk in **A**) were labelled with MitoTracker Red and co-cultured with unlabelled primary human myocytes (hMYO) for 4 hours followed by staining with phalloidin-Alexa Fluor 488. Nuclei were stained with DAPI. Mitochondrial transfer (orange-red, MITO) from hM1-Mac (yellow in merged images **A, B**) to hMYO (green, shown in **A, C**) occurs through tunneling nanotubes (TNTs) (marked with yellow arrows in **A** and **C**). **A** = merged image; **B** = red and blue channels showing mitochondrial transport through TNTs; **C** = green and blue channel showing F-actin; **D** = blue channel showing DAPI. Scale bar = 25 µm.

To examine the TNT formation and mitochondrial transport between M1-Macs and MYO in real time, we performed time-lapse imaging of the primary hM1-Macs and hMYOs (**Video 1**, **Figure 3A**). Prior to co-culture, mitochondria within hM1-Macs were labeled with MitoTracker (far-red), and their plasma membrane was stained with a red dye to visualize TNT projections. hMYOs were stained with wheat-germ agglutinin (WGA)-AlexaFluor-488 (green). The time-lapse imaging revealed that: **1)** TNTs between M1-Macs and MYO are transient in nature, repeatedly forming and disappearing during time-lapse imaging (**Video 1**, **Figure 3A**); **2)** TNTs originate from hM1-Macs and formed connections with hMYO as evidenced by the TNT structures labelled with the red-plasma membrane dye, (**Figure 3B-D**); **3)** mitochondria labeled within hM1-Macs are transported through TNTs to hMYOs, observed as yellow punctate staining accumulating within hMYOs over time (Video 1, **Figure 3A**); **4)** labeled mitochondria are rapidly shared among neighboring hMYOs following their transfer from hM1-Macs, suggesting that hMYOs also can communicate with each other via TNTs (**Supplemental Video 3, Supplemental Figure 5**); **5)** no TNT connections or mitochondrial transfer was observed between monocyte-derived hM2-Macs and hMYOs (**Supplemental Figure 6**). This finding demonstrate that TNT formation is specific to M1-Macs, but not to M2-Macs.

**Figure 3:**
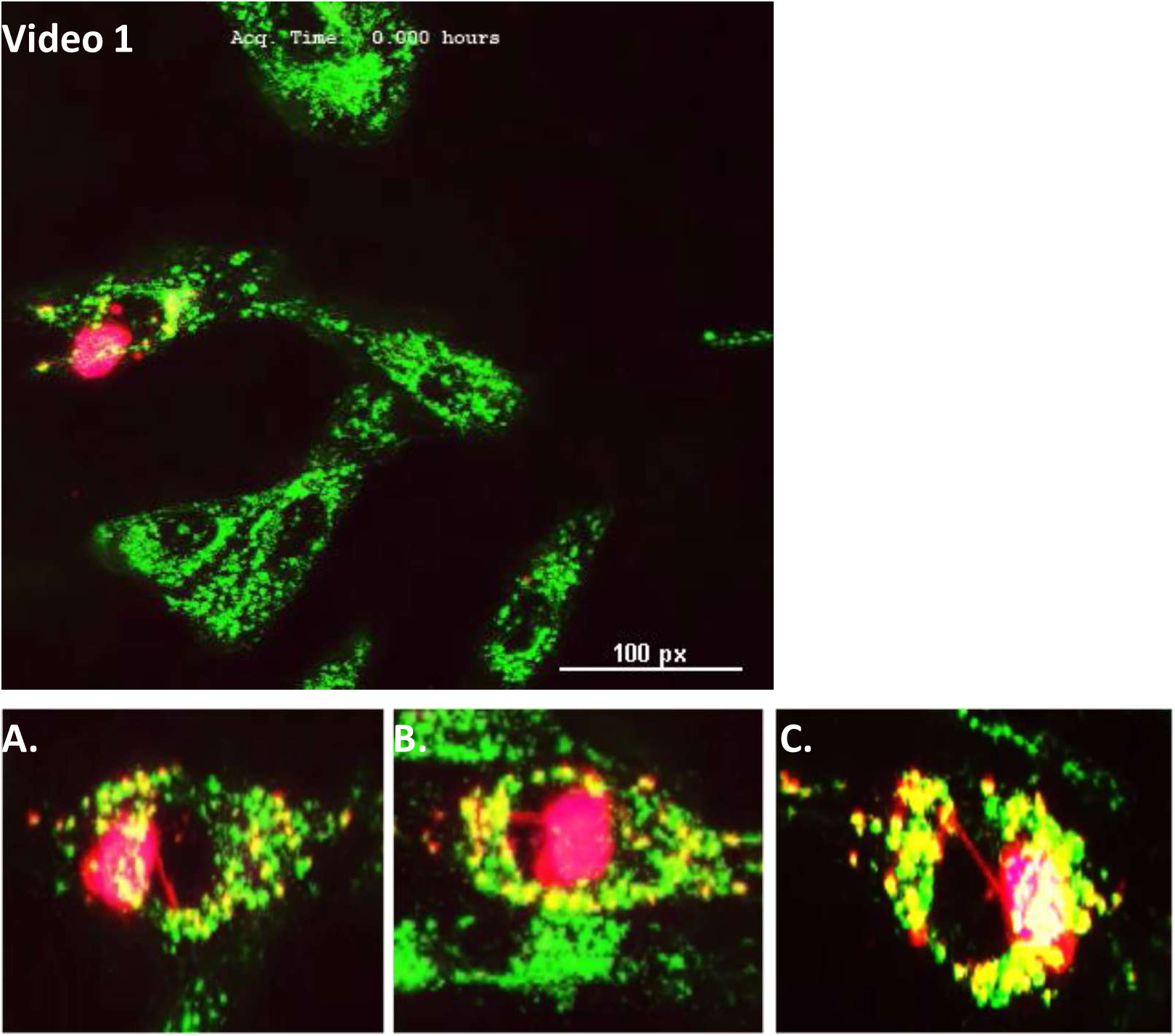
Time-lapse imaging of tunneling nanotube formation and intercellular mitochondrial transfer between M1 macrophage and myocytes. Human PBMC-derived M1 macrophages (hM1 Macs; labelled with MitoTracker, shown in pink, and a plasma membrane marker, red) were co-cultured with wheat germ agglutinin-labelled human myocytes (hMYO). Time-lapse imaging was performed over ∼15 hours. **Video 1)** Representative time-lapse video capturing dynamic TNT formation and mitochondrial movement. **A–C)** Selected time-point images illustrating various stages of TNT formation and mitochondrial transfer between hM1-Macs and hMYO. Scale bar = 100 pixels = 100 µm.

### Mitochondrial transfer via TNT occurs in a unidirectional manner from M1-Mac to MYO

Next, we investigated whether mitochondrial transfer occurs unidirectionally from M1-Macs to MYOs or as a bidirectional exchange between Macs and MYOs. We utilized a transgenic mitodendra2-C57-BL/6 (mitoDendra) mouse model, which ubiquitously expresses mitochondria targeted Dendra2 which without photoconversion emits green fluorescence. Peripheral monocytes were isolated from pregnant mitoDendra mice on GD15 and differentiated *in vitro* into M1-Macs (mitoDendra-M1-Macs). Primary murine myocytes (mMYOs) were isolated from pregnant (GD15) wild-type C57BL/6 (WT) mice, stained with the F-actin probe SPY650 (red) and co-cultured with mitoDendra-M1-Macs, followed by time-lapse imaging to assess mitochondrial transfer. Consistent with the human M1-Mac-MYO co-culture model (using fluorescent MitoTracker dye), this murine co-culture model also demonstrated the transfer of mitochondria from mitoDendra-M1-Macs to mMYOs **(Video 2, Figure 4, Supplemental Video 4, 5).**

**Figure 4:**
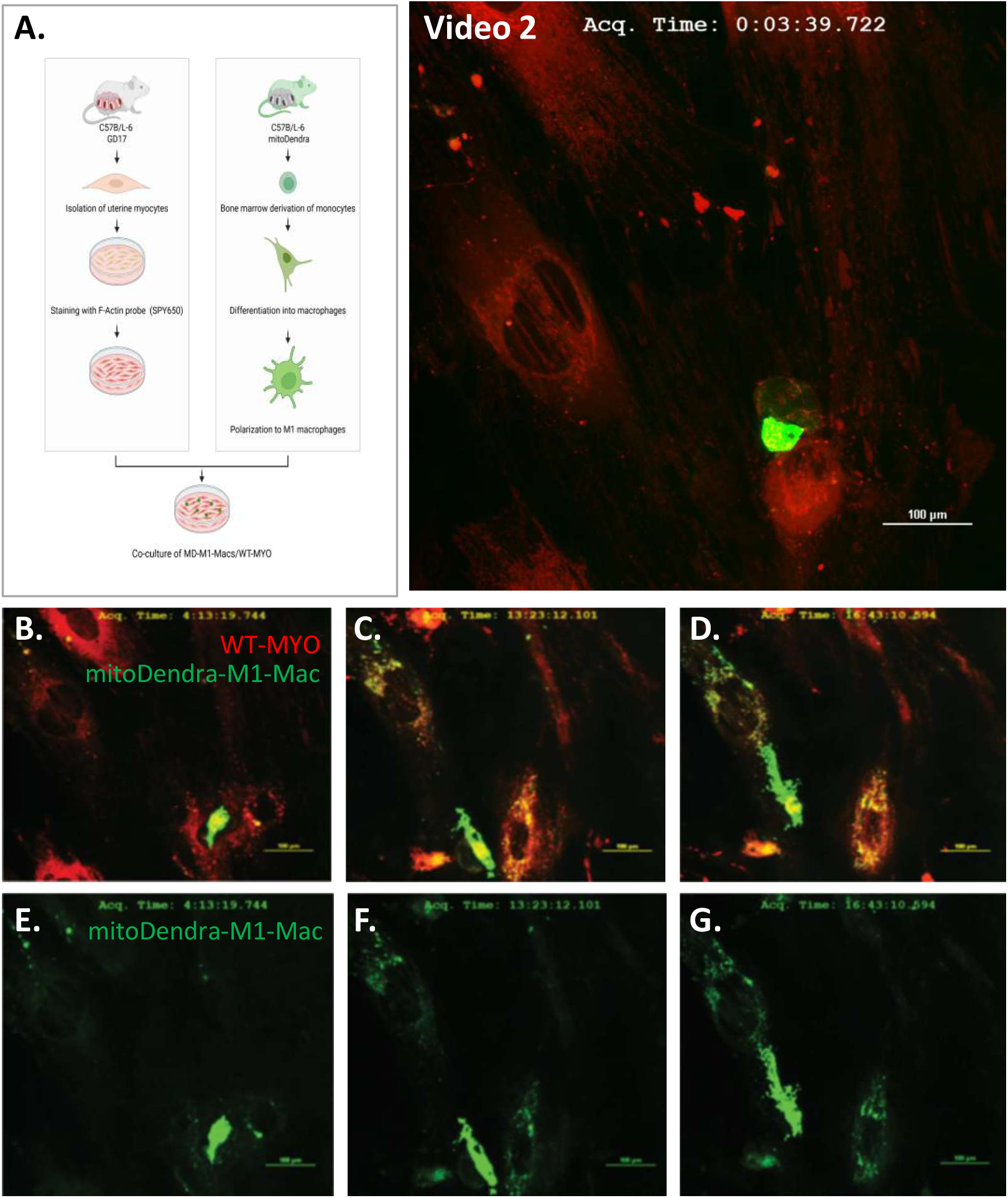
Mitochondrial transfer from M1 macrophage to myocytes is unidirectional: Time lapse imaging of primary murine mitoDendra-M1-macrophages (mitoDendra-M1-Macs) with dendra-labelled mitochondria and wild type myocytes (WT-mMYO) in co-culture. **A)** Schematic representation of experimental design. mM1-Macs were derived from the peripheral blood monocytes from pregnant mitodendra2-CD57B/L-6 (mitoDendra) mice, which ubiquitously express Dendra fluorescence in mitochondria, on gestational day (GD) 15. Primary murine myocytes were isolated from wild type CD57B/L-6 (WT) mice on GD15 and stained with the F-actin probe; SPY650. Co-culture was established between the mitoDendra-M1-Macs and WT-mMYO, and time-lapse imaging was performed overnight. The transfer of dendra-labelled (green fluorescent) mitochondria from mitoDendra-M1-Macs to WT-mMYO was observed as shown in **Video 2**. **(B–G)** Representative time-lapse images at the indicated time points. Scale bar = 100 µm.

In the reciprocal experimental setup, M1-Macs were derived from WT mice and labelled with a far-red plasma membrane dye, while primary mMYOs were isolated from mitoDendra mice (mitoDendra-mMYO). No mitochondrial transfer from mitoDendra-mMYO to WT-M1-Macs was observed in these co-cultures indicating that mitochondrial transfer occurs in a unidirectional manner from M1-Macs to MYOs **(Video 3, Figure 5)**.

**Figure 5:**
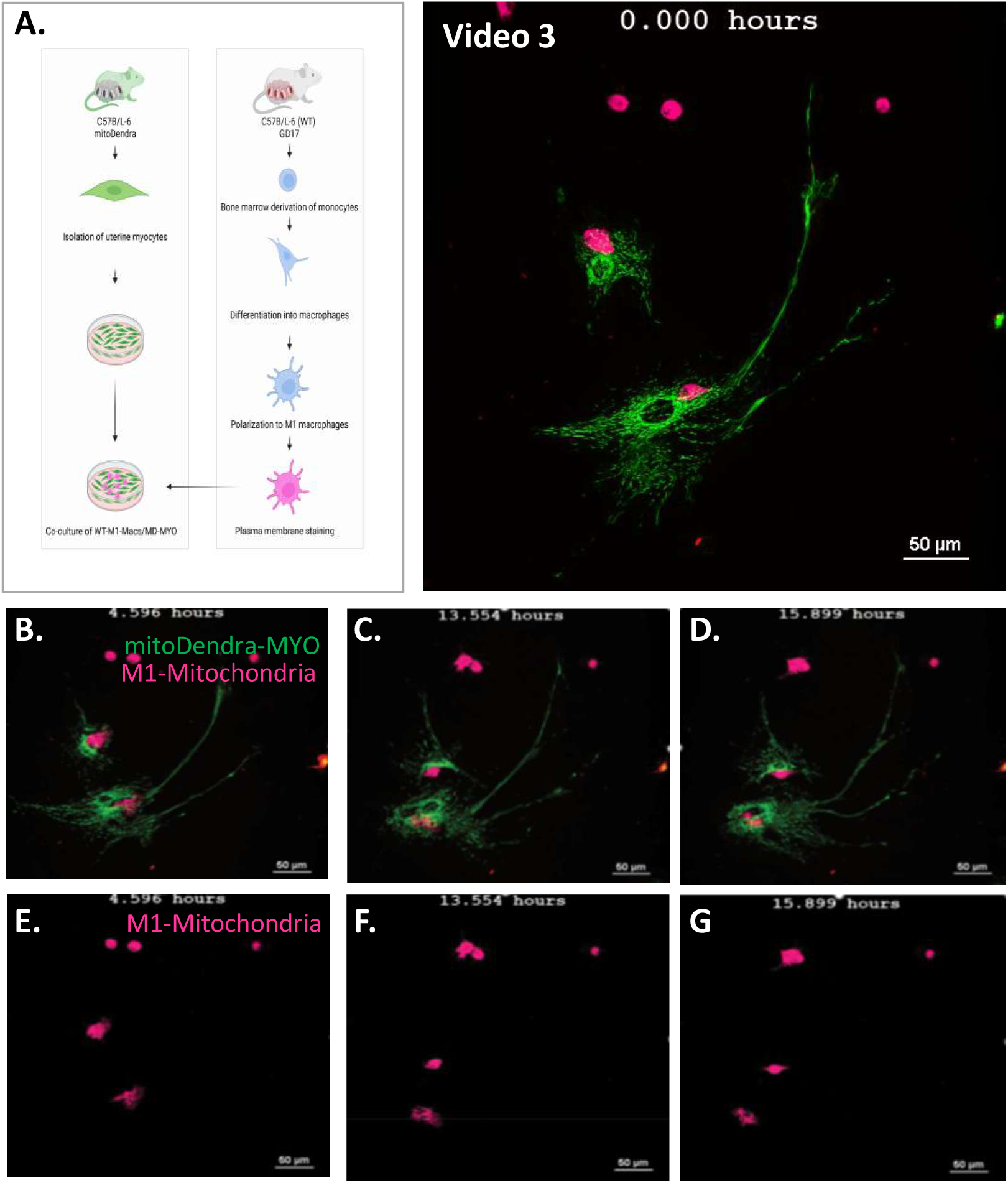
Mitochondrial transfer from M1 macrophage to myocytes is unidirectional: Time lapse imaging of primary murine WT-M1-macrophages (WT-mM1-Macs) and mitodendra2-CD57-BL/6 uterine myocytes (mitoDendra-MYO) with dendra-labelled mitochondria in co-culture. **A)** Schematic representation of experimental design. Primary murine myometrial cells were isolated from mitoDendra2-C57BL/6 transgenic mice (mitoDendra-mMYO), which ubiquitously express the Dendra fluorescent protein targeted to mitochondria. M1 macrophages were differentiated from peripheral blood monocytes isolated from wild-type C57BL/6 mice (WT-mM1-Macs), labelled with Vybra DiD Cell-Labeling Dye, and co-cultured with mitoDendra-mMYO cells. Time-lapse imaging was performed overnight to assess mitochondrial transfer. No transfer of Dendra-labelled (green fluorescent) mitochondria from mitoDendra-mMYO cells to WT-mM1-Macs was observed, as demonstrated in **Video 3**. **(B–G)** Representative time-lapse images at the indicated time points confirm the absence of mitochondrial transfer from myocytes to macrophages, supporting the unidirectional nature of mitochondrial transfer from M1 macrophages to myocytes.

### Myocytes modulate TNT formation by M1-Macs

TNF alpha-induced protein 2 (TNFAIP2, also known as M-Sec) is a well-established marker of TNTs ^29^. We first examined the expression of TNFAIP2 in co-cultures established between human myometrial cells and monocytes (MYO/M), non-polarized Macs (MYO/M0-Macs), MYO/M1-Macs and MYO/M2-Macs. TNFAIP2 protein was detected in MYO monoculture and all co-cultures tested (**Figure 6A**). Importantly, TNFAIP2 protein levels were significantly higher in the MYO/M1-Mac co-cultures compared all other co-cultures assessed. When TNFAIP2 protein levels were compared in monocultures of these cell types, M1-Macs showed highest levels of TNFAIP2 protein (**Figure 6B**).

**Figure 6:**
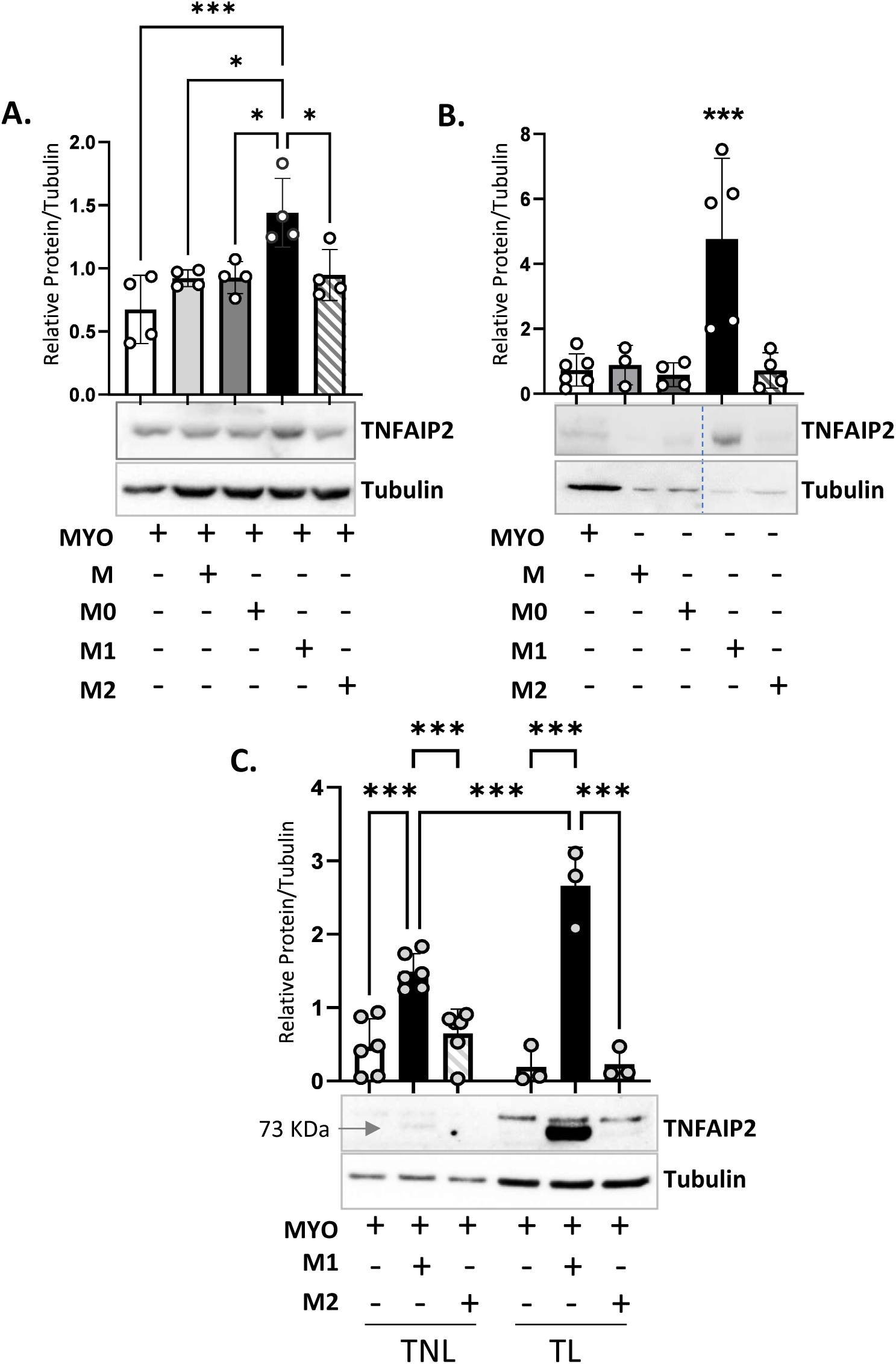
M1 macrophages express high levels of the TNT marker M-Sec/TNFAIP2. Representative western blot images and densitometric analysis of M-Sec/TNFAIP2 protein. Comparative levels of M-Sec/TNFAIP2 are shown in; **A)** hTERT-HM^A/B^ myocytes (MYO) in monoculture versus when they are co-cultured with THP-1 monocytes (M) or THP-1 derived macrophages (M0, M1, M2) for 24 hours, **B)** MYO, M, M0, M1 and M2 in monoculture, and **C)** primary human MYO derived from term not in labor (TNL) versus term in labor (TL) women, in monoculture and when co-cultured with PBMC derived M1 or M2 macrophages. Graphs show TNFAIP2 normalized with Tubulin protein (loading control). Data are presented as mean ± SD (N=3-6). Statistical comparisons were made between the MYO in monoculture with other groups. Statistical significance was determined by One-way ANOVA followed by Šídák’s multiple comparisons test (A), Dunnett’s multiple comparisons test (B), or Two-way ANOVA with Šídák’s multiple comparisons test (C). ‘***’ denotes p ≤ 0.001 and ‘*’ p ≤ 0.033. MYO (white bars), M-monocytes (light grey bars), non-polarized macrophages (M0, dark grey bars); M1 polarized macrophages (M1, black bars); M2 polarized macrophages (M2, striped bars).

Notably, when primary hM1-Macs and hM2-Macs were co-cultured with hMYO derived from term non-labour (TNL) vs term labour (TL) myometrium, hM1-Macs co-cultured with TL-hMYO showed significantly higher levels of TNT marker compared to hM1-Macs co-cultured with TNL-hMYO (**Figure 6C**). These findings suggest that MYOs may promote TNT formation from M1-Macs through paracrine signaling during human labour.

### M1-Macs induce functional and intracellular progesterone withdrawal in the myocytes

To investigate the influence of Macs on intracellular and functional P4 withdrawal in MYO, THP-1 (M) derived Macs (M0) were polarized into M1-Macs or M2-Macs and co-cultured with MYOs (hTERT-HM^A/B^ cell line, Mac:MYO ratio 1:10). Our results showed that M1-Macs significantly upregulate the expression of 20α-HSD (**Figure 7A**), induce phosphorylation of PR-A at the S^344/345^ residues (**Figure 7B**), decrease PR-B, increase PR-A/PR-B ratio (**Figure 7C**) and elevate levels of connexin 43 (CX43) (**Figure 7D**). Notably, these effects were not evident when MYOs were co-cultured with monocytes, M0 or M2-Macs.

**Figure 7:**
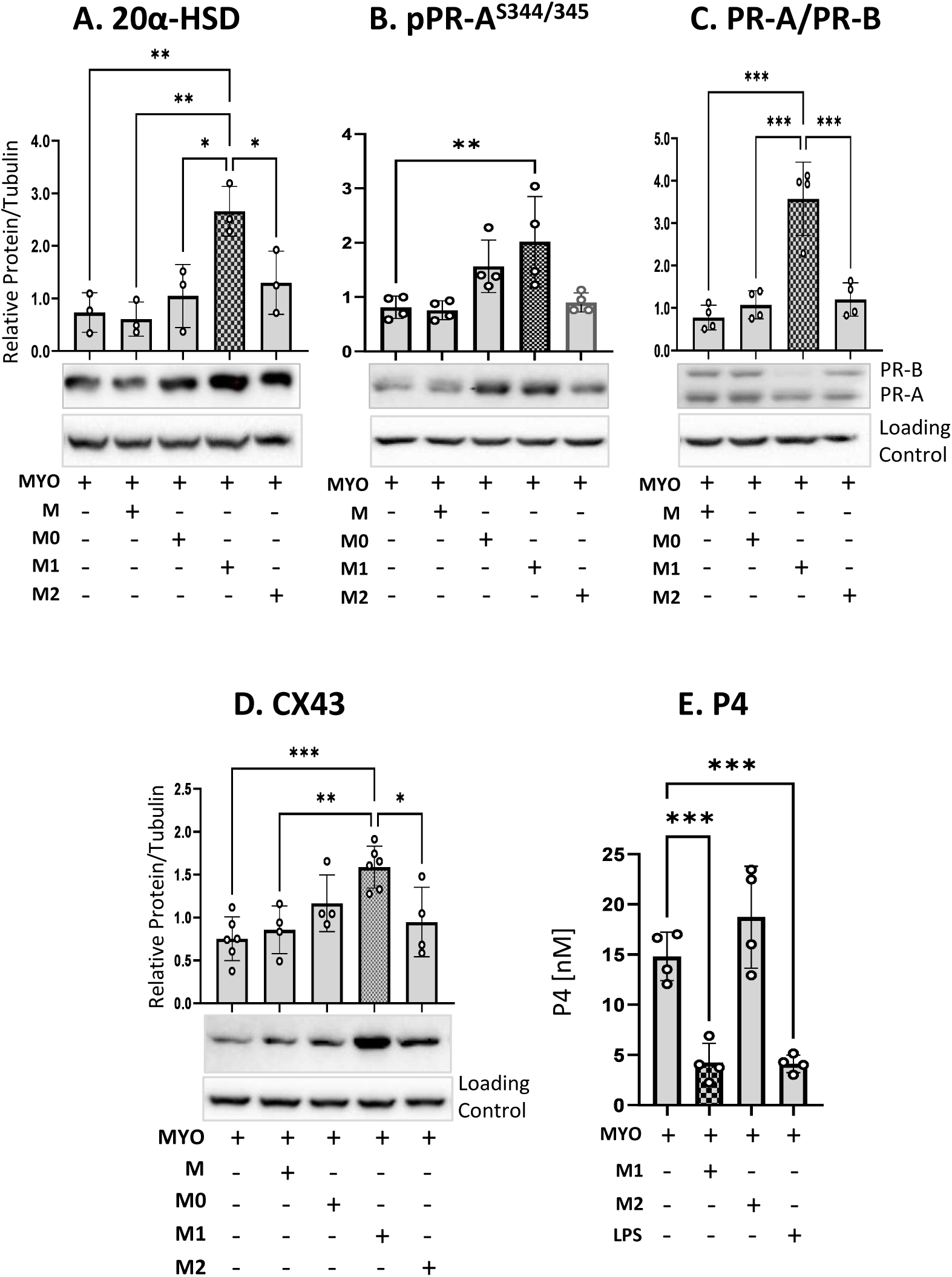
M1 macrophages induce 20α-HSD-mediated intracellular and PR-A-phosphorylation-mediated functional progesterone withdrawal in myocytes. Representative western blot images and densitometric analysis of Human monocytic cell line; THP1 (M) was differentiated into Macrophages (M0), polarized into M1 and M2 phenotypes and co-cultured with hTERT-HM^A/B^ cells (MYO), in the ratio of 1:10 (M1:MYO) in the presence of 25 ng/ml DOX, 100 nM GSL-1 and 100 nM progesterone (P4). Graphs show comparative levels of 20α-HSD (**A**), phosphorylation of PR-A at S344/345 (**B**), levels of total PR-A and PR-B (**C**), and CX43 (**D**). Graphs show protein levels relative to Tubulin. Data are presented as mean ± SD (N=3-4 independent experiments). **E) M1 macrophages reduce progesterone levels in myocytes.** hTERT-HM^A/B^ cells (MYO), induced to express PR-A/PR-B were cultured alone with/out LPS (100 ng/mL) (positive control), or co-cultured with THP-1 derived M1 or M2 macrophages for 18 hours in the presence of progesterone. Progesterone levels (P4) were assessed in the culture medium. Graph shows mean ± SD (N=3-4 independent experiments). Significant comparisons are denoted by ‘***’ p≤0.001**’, p≤0.002 and ‘*’ p≤0.033’ MYO (white bars), M-monocytes (light grey bars), non-polarized macrophages (M0, dark grey bars); M1 polarized macrophages (M1, black bars); M2 polarized macrophages (M2, striped bars).

To elucidate the impact of elevated 20α-HSD expression on P4 metabolism, we measured P4 levels in the conditioned media from of hM1-Mac/hMYO and hM2-Macs/hMYO co-cultures using ELISA. hM1-Macs/hMYO co-cultures caused a significant reduction of P4 levels, compared to hM2-Mac/hMYO co-cultures and hMYO monocultures (**Figure 7E**). The reduction in P4 levels mediated by M1-Macs was comparable to the decrease observed following LPS treatment of MYOs. LPS served as a positive control, since it reduces P4 levels via induction of 20α-HSD (as shown earlier ^6^).

### M1-Mac/MYO co-culture increases total ATP levels in myocytes

As shown in **Figure 3**, hM1-Macs/hMYO co-culture results in transfer of mitochondria from M1-Macs to MYO, which could change cellular energetics. To evaluate this, we measured the adenosine triphosphate (ATP) levels in individual cell populations; hMYO, hM1-Macs and hM2-Macs in monoculture, and in hM1-Macs/hMYO or hM2-Macs/hMYO co-cultures (Macs:MYO ratio1:10). A significant increase was observed in total ATP content in hM1-Macs/hMYOs co-culture as compared to monocultures or hM2-Macs/hMYO co-culture (**Figure 8A**). When Macs were pre-treated with metformin (1 mM), a well-characterized mitochondrial function inhibitor ^30^, for one hour prior to co-culture, no increase in ATP content in hM1-Mac/hMYO co-cultures was detected, indicating that the ATP induction in MYO is due to the mitochondrial transfer from M1-Macs to MYO (**Figure 8B**). To support this conclusion, fluorescently labelled hM1-Macs were pre-treated with metformin (1 mM) for one hour prior to co-culture with hMYO; our data show a complete blockade of TNT formation and mitochondrial transfer from hM1-Mac to hMYO in the presence of metformin (**Figure 8C**).

**Figure 8:**
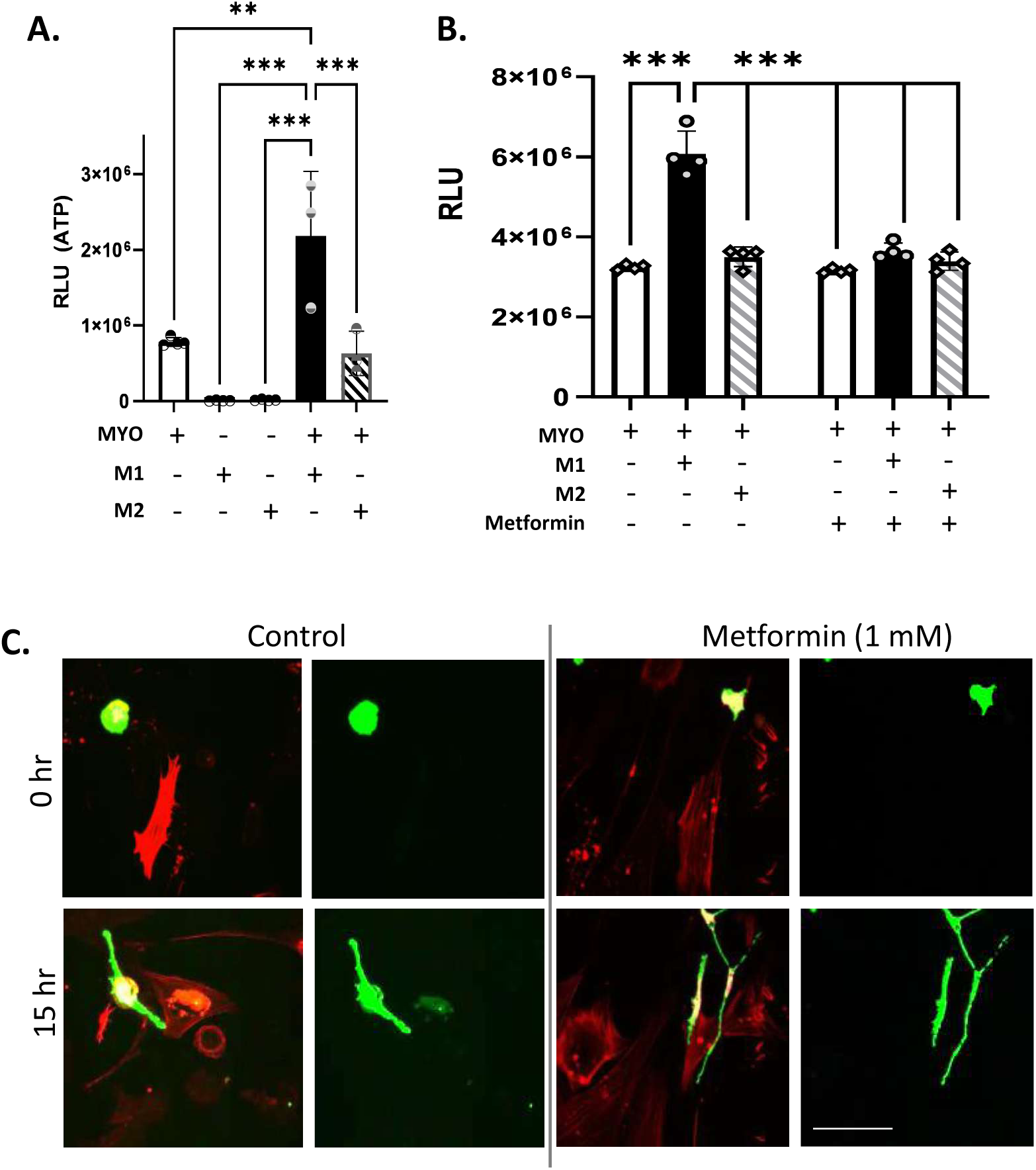
Metformin blocks transcellular transfer of mitochondria from M1 macrophages to myocytes. A) M1-Mac/MYO co-cultures have higher ATP content compared to monocultures. ATP levels were assessed in 100,000 MYO, 10,000 M1 and M2 cells in monoculture and when they were co-cultured for 24 hrs. Graph shows mean ± SD (N=3 independent experiments). Statistical significance was determined between MYO monoculture *vs* other groups by one-way ANOVA followed by Dunnett’s multiple comparison test. Significant comparisons are denoted by ‘***’ p≤0.001, ‘**’ p≤0.002. **B)** THP-1 derived M1 and M2 Macs were pretreated with Metformin (1 mM) or control for one hour and co-cultured with hTERT-HM^A/B^ cells (hMYO) in a ratio of 1:10 for 24 hours. Total ATP level was measured. Graph shows mean ± SD (N=4 independent experiments). Statistical significance was determined between groups by two-way ANOVA followed by Tukey’s multiple comparison test. Significant comparisons are denoted by ‘***’ p≤0.001. MYO (white bars), M1 polarized macrophages (M1, black bars); M2 polarized macrophages (M2, striped bars). **C)** mitoDendra-M1-Macs were derived from the peripheral blood monocytes from mitodendra2-CD57B/L-6 (mitoDendra) mice which ubiquitously express Dendra fluorescence in mitochondria. Primary murine myocytes (mMYO) were isolated from wild type CD57B/L-6 (WT) and stained with the F-actin probe; SPY650 (shown in red). mitoDendra-M1-Macs were pretreated with Metformin (1 mM) or control for one hour and co-cultured with WT-mMYO. Time lapse imaging was performed overnight. Snapshots of 0 and 15 hours are shown. Scale bar = 100 µm.

### Mitochondria from M1 Macrophages induce 20α-HSD Levels in Myocytes

As the expression of 20α-HSD was significantly upregulated in human M1-Macs/MYO co-cultures (**Figure 7A**), we hypothesized that this effect is due to mitochondrial transfer from M1-Macs to MYO. To validate our methodology, we first demonstrated that isolated (naked’) mitochondria are taken up by the MYOs when added to the culture medium. We isolated Dendra^+ve^ mitochondria from mM1-Macs obtained from mitoDendra mice and co-cultured them with mMYOs for 24 hours. The internalization of Dendra^+ve^ mitochondria by mMYOs was confirmed by fluorescence imaging (**Supplemental Figure 2**). To assess the role of mitochondrial transfer on MYOs, human M1-mitochondria were isolated from THP-1-derived M1-Macs and added to the human MYOs monoculture. Mitochondrial uptake led to a significant induction of 20α-HSD levels in MYOs (**Figure 9B**), comparable to the increase observed in M1-Macs/MYOs co-culture (**Figure 7A**). However, in contrast to the M1-Mac/MYO co-culture, PR-A phosphorylation remained unaffected by the mitochondrial uptake (**Figure 9C**). Pre-treatment of M1-Macs with Metformin (effective dose = 1mM) before transplantation led to a marked downregulation of 20α-HSD (**Figure 9D**) and CX43 protein levels (**Figure 9E**) in MYOs, while PR-A phosphorylation levels remained stable (**Figure 9F**). These findings indicate that mitochondrial transfer to MYOs contributes to the induction of 20α-HSD, and subsequent induction of CX43 expression, while functional P4 withdrawal (and PR-A dominance) is achieved independent of the mitochondrial transfer.

**Figure 9:**
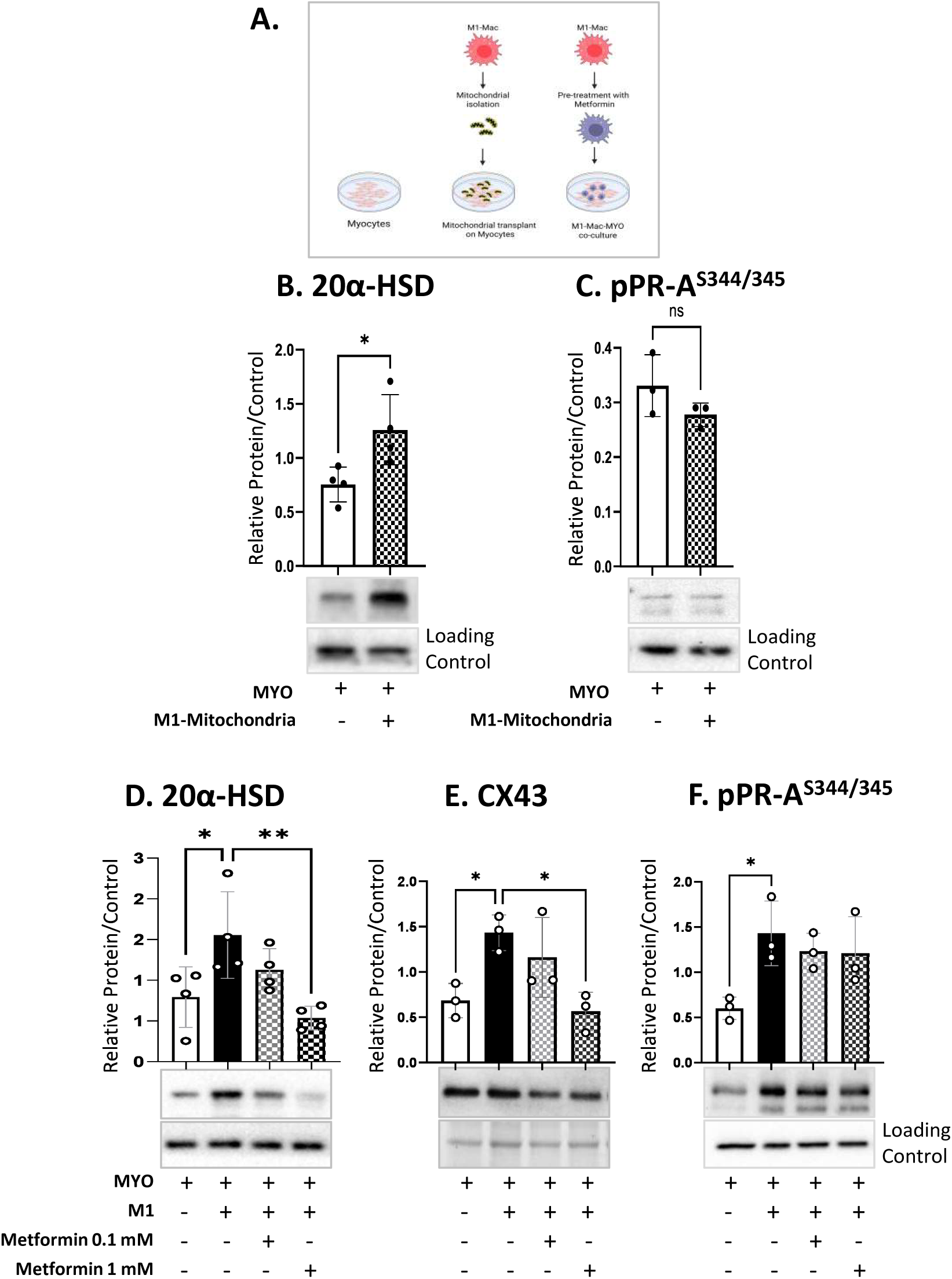
Mitochondrial transfer from M1 macrophages to myocytes induces 20α-HSD levels in myocytes. **A)** Schematic representation of mitochondrial transplant experiment. **B, C)** Mitochondria were isolated from the THP-1-derived M1-Macs (hM1-Mitochondria). Naked mitochondria were transplanted on the hTERT-HM^A/B^ cells (hMYO), for 24 hours and levels of 20α-HSD (B) and pPRA^S344/345^ (C) were determined via western blotting. **D-F)** THP-1-derived hM1-Macs were pretreated with Metformin (0.1 mM or 1 mM) or control for 1 hour and then co-cultured with hMYO for 24 hours. Levels of 20α-HSD (**D**), pPR-A^S344/345^ (**F**) and CX43 (**E**) was determined via western blotting. Graphs represent protein levels normalized to the appropriate loading controls: Tubulin for panels B–D, total protein (stain-free gel) for panel E, and ERK2 for panel F. Data are presented as mean ± SD (N=3-4 independent experiments). Statistical significance was determined between two groups by student’s ttest (**B, C**) and between more than two groups by one-way ANOVA followed by Dunnett’s multiple comparison test (M1 vs other groups, **D-F)**. Significant comparisons are denoted by, ‘**’ p≤0.002 and ‘*’ p≤0.033. MYO (white bars), M1 polarized macrophages (M1, black bars); Metformin (checkered bars).

### *In vivo* evidence of monocyte infiltration, differentiation to M1-Macs and intercellular mitochondrial transfer in the pregnant uterus

To establish the physiological relevance of the TNT-mediated mitochondrial transfer from M1-Macs to MYOs in relation to labor onset, we used the mitoDendra mouse model described above. First, we isolated Dendra2^+ve^ monocytes from the peripheral blood of pregnant mitoDendra mice on gestational day 15 (GD15). These monocytes were adoptively transferred into GD15 pregnant recipient WT C57BL/6 mice via tail vein injection (monocyte numbers ranging from 40,000 to 100,000 cells/dam). Uteri from injected WT mice were collected at 24-hour intervals throughout gestation (GD16, 17, 18, 19/TNL), TL and postpartum (PP) and the presence of the Dendra2^+ve^ cells analyzed (**Figure 10A**).

**Figure 10:**
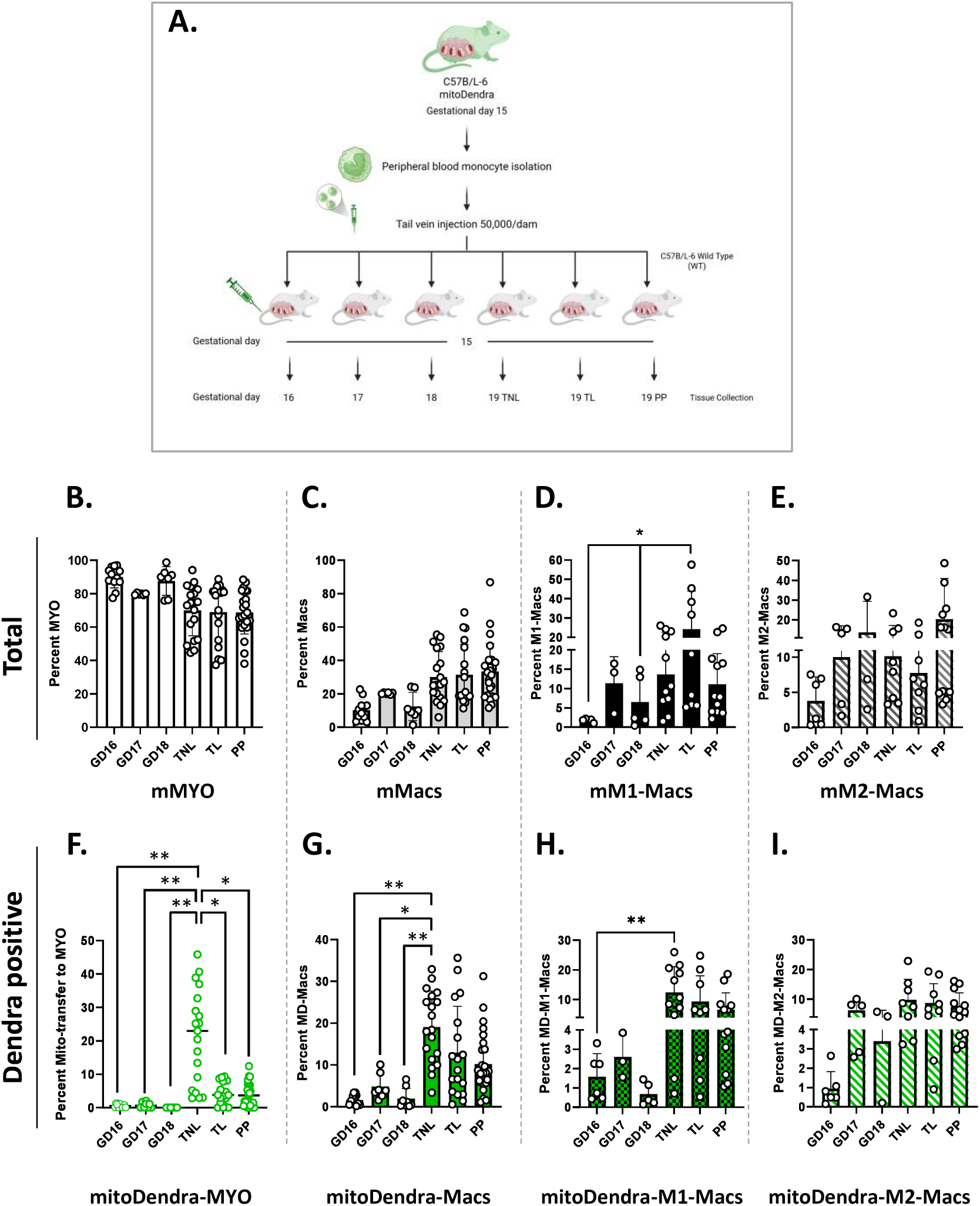
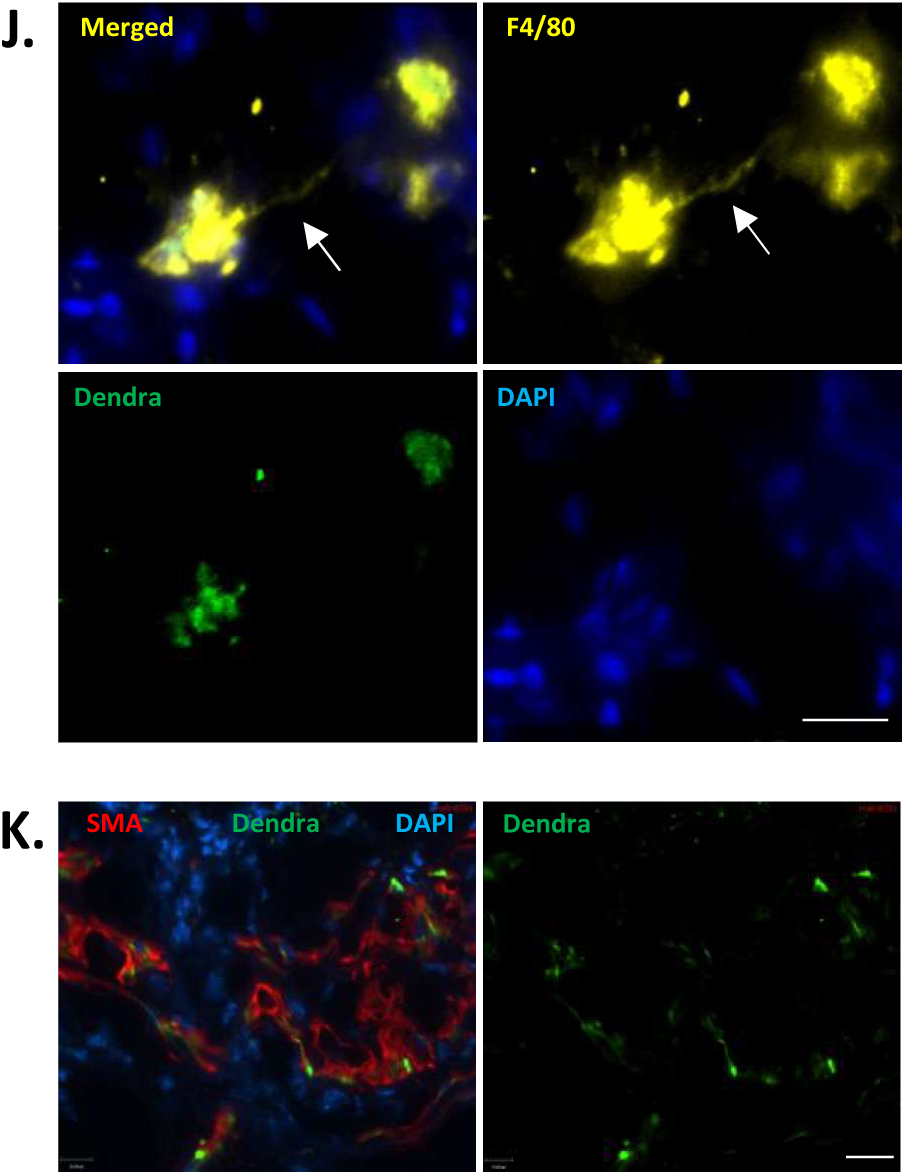
In vivo evidence of monocyte infiltration, differentiation to M1 macrophages and transcellular mitochondrial transfer in the pregnant uterus. **A)** Schematic representation of the experimental design for adoptive transfer of monocytes. Illustration created in https://BioRender.com. Peripheral blood monocytes were isolated from pregnant C57BL/6 MitoDendra2 (mitoDendra) mice on gestational day (GD) 15. Adoptive transfer (50,000–100,000 monocytes per dam) was performed on the same day into pregnant wild type C57BL/6 wild-type (WT) recipient mice via tail vein injection (N = 3 per gestational day). Uterine tissues were collected at 24-hour intervals post-injection. **B-I)** Quantitative analysis of myometrium by QuPath (7.0). **B-E)** Graphs represent the total population of indicated cell types (dendra^-ve^ plus dendra^+ve^) myocytes (mMYO, white bars), macrophages (mMacs, grey bars), pro-inflammatory mM1-Macs (black bars), and anti-inflammatory mM2-Macs (stripped bars). **F-I)** Graphs represent infiltrated population of mMacs (dendra^+ve^): mitoDendra-Macs (**G**), mitoDendra-M1-Macs (**H**) and mitoDendra-M2-Macs (**I**), **F)** Shows dendra ^+ve^ Myocytes (mitoDendra-MYO) representing MYO with mitochondria acquired from infiltered Macs. Graphs show percent cells, calculated from the total number of cells detected per region of interest per tissue section. Data are presented as mean ± SD (N=3 mice per gestational day). Statistical significance was determined by one-wray ANOVA followed by dukey’s multiple comparison test (all groups compared with each other). Significant comparisons are denoted by ‘**’ p≤0.002, ‘*’ p≤0.033. **J)** Capture of TNT formation in-vivo (white arrow) originating from mitodendra^+ve^ (green) and F4/80^+ve^ Mac (yellow) in term myometrium tissue. DAPI represents MYOs presence in the surrounding. **K**) Transfer of mitoDendra-derived mitochondria to MYO is shown *in-vivo* at term. Smooth muscle actin marks MYOs (red) and mitoDendra-derived mitochondria (green). Scale bar = 20 µm.

Validated marker antibodies were used to identify specific cell populations in murine uterus, including mMacs (F4/80^+ve^), mM1-Macs (F4/80^+ve^CD38^+ve^), mM2-Macs (F4/80^+ve^Egr2^+ve^), and mMYO (SMA^+ve^) as described in ^24^. Dendra positive-Mitochondria were detected using PLA amplification with dendra specific antibody and +/-probes targeting the same antibody. Multiplexed immunostaining was performed on adjacent uterine sections to detect fluorescent Dendra^+ve^ cells in SMA^+ve^ myometrium. Quantification showed the SMA^+ve^ mMYO population remained unchanged (**Figure 10B**), while the total F4/80^+ve^ mMacs population trended higher with advancing gestation without attaining statistical difference (**Figure 10C**). Importantly, a significant increase in the total number of mM1-Macs (which is a combination of resident mMacs plus mitoDendra-Macs differentiated from infiltrated Dendra^+ve^ monocyte) was observed during labor (**Figure 10D**). Dendra^+ve^ mMacs were detectable in the myometrium throughout gestation, starting from GD16 (24 hours post injection). The number of Dendra^+ve^ mMacs and Dendra^+ve^ mM1-Macs significantly increased on GD19 before the onset of labor/TNL (**Figure 10G, H**). Dendra positivity was also observed in the mM2-Macs throughout gestation, with no significant change in their proportion (**Figure 10I**). Notably, a significant increase in the number of Dendra^+ve^ mMYO was detected on GD19/TNL which later declined as the labor progress and postpartum (mitoDendra-MYO, **Figure 10F**, **Supplemental Figure 3**), indicating that the transcellular mitochondrial transfer primarily occurs at term prior to labor onset.

Collectively these in-vivo data suggest that Dendra^+ve^ monocytes infiltrate the uterus within 24 hours of adoptive transfer. These monocytes differentiate to Macs as evident by co-expression of Mac-specific marker F4/80 and Dendra. The infiltered Macs polarize to both M1 or M2-Macs during GD16-18, and the mitoDendra-M1-Macs population increases on GD19/TNL before labor onset, while mitoDendra-M2-Macs numbers remain constant during late gestation, TL and PP. Importantly, direct *in vivo* evidence of TNT formation was observed in murine myometrial tissue from a term pregnant dam (**Figure 10J**). Dendra positivity in mMYO was determined by the detection of dendra in smooth muscle actin (SMA) positive cells (**Figure 10K).**

## Discussion

The molecular mechanisms driving the transition of the pregnant myometrium from quiescence to contractility during labor remains poorly understood. A substantial and growing body of evidence implicates the immune system as key contributor of both term and preterm labor. The present study demonstrates, for the first time, a novel mechanism of communication between the maternal immune and reproductive systems that triggers labor (via P4 withdrawal) and supports the energetic demands of uterine contractions.

TNT-mediated open-channel connections between M1-Macs and MYO in the pregnant uterus facilitate intercellular cargo transfer. This interaction leads to myometrial P4 withdrawal, inducing intercellular connectivity via CX43, and provides supplemental energy to support labor. The temporal pattern of these events suggests a mediating role of this interaction in the initiation of labor. Among the two classical Mac subtypes (M1 and M2), only M1-Macs show the capability to form rapid and direct TNT connections with MYOs, as evidenced by the high expression of TNT marker TNAIP2 in M1-Macs and mitochondrial transfer exclusively from M1-Macs to MYO. In contrast, M2-Macs do not exhibit these features (**Supplemental Figure 6**). Evidence suggests that M2 macrophages can form TNT connections in other tissues ^31^; however, in our study, we did not observe TNT formation with uterine myocytes by M2-Macs derived from PBMCs or THP-1 monocytes. Recent studies have reported the presence of TNT-mediated communication among myeloid cells, including Macs, that support intercellular signaling, pathogen transfer, and the coordination of immune responses ^32^. Macs have been shown to form both homotypic (Mac–Mac) and heterotypic (Mac–other cell types) TNT connections with epithelial, stromal, HIV-infected and tumor cells ^32, 33, 34^ however, to our knowledge, this is the first evidence demonstrating TNT formation between M1-Macs and uterine MYOs, with the functional consequence of this interaction linked to the onset of labor.

The exclusive observation of unidirectional cargo transfer from M1-Macs to MYO is particularly intriguing and suggests that this process may be tightly regulated, potentially driven by the MYOs, as the primary beneficiary in unidirectional transfer is generally the recipient cell ^35^. In the context of labor, the ability of M1-Macs to rapidly form TNT connections with both nearby and distant MYOs suggests that even a small population of activated M1-Macs can efficiently propagate signals and provide metabolic support across the tissue within a short timeframe. This concept is further supported by the observed increase in MYO-MYO connectivity in the presence of M1-Macs, as indicated by the distribution of M1-derived cargo among myocytes that are not directly connected to Macs via TNTs. (**S-Video 3**, **Supplemental Figure 5**). Interestingly, the TNT formation between M1-Mac/MYO and the enhanced connectivity between MYOs was only evident in the presence of P4, suggesting that P4 plays an important role in facilitating M1-Macs/MYO communication and cargo transfer (**Supplemental Figure 4**).

Mitochondria are unlikely to be the only cargo transferred during Mac-MYO crosstalk. To confirm this, we performed time-lapse imaging of hM1-Macs labelled with MitoTracker Red (red, appearing yellow in merged image) and the plasma membrane co-labelled with wheat germ agglutinin (WGA, green). Immunofluorescence imaging confirmed the presence of two distinct cargo types from M1-Macs to MYOs: mitochondrial cargo, identified by its yellow fluorescence in merged images, and macrophage plasma membrane-associated cargo, identified by green fluorescence. The fluorescence signals visually distinguish these distinct cargo populations (**Supplemental Figure 7**).

The functional consequence of M1-Macs/MYO communication resulted in three key cellular outcomes in MYOs: **1)** intracellular P4 withdrawal through 20α-HSD mediated metabolism, accompanied by functional P4 withdrawal as reflected by increased PR-A phosphorylation and elevated PR-A/PR-B ratio; **2)** a significant increase in cellular ATP levels, and 3) activation of hMYO as indicated by increased expression of the contraction-associated protein, CX43 level. Transfer of mitochondria isolated from hM1-Macs to hMYO only partially mimicked the M1-Macs-induced P4 withdrawal. Although mitochondrial transfer was able to induce 20α-HSD expression, it did not alter PR-A phosphorylation or the PR-A/PR-B ratio. These findings suggest that additional M1-Mac-derived factors contribute to the modulation of PR signaling, potentially including IL1α or IL-1β cytokines, as previously reported ^7, 36^.

The increased ATP levels in MYOs suggest that functional mitochondria are transferred from M1-Macs to MYOs and can be utilized by MYOs as an energy resource to facilitate the process of labor. The increase of ATP levels in M1-Mac/MYO co-cultures is attributed to mitochondrial transfer from M1-Macs to MYO. This interpretation is supported by the experiments using pretreatment of M1-Macs with metformin, which inhibited both mitochondrial transfer and the associated rise in ATP levels in myocytes. The inhibitory effect of metformin on transcellular mitochondrial transfer may be explained by the energy dependence of intracellular mitochondrial trafficking. Mitochondrial movement relies on interactions with cytoskeletal and motor protein complexes, including kinesin/Miro1, along actin filaments within TNTs. Transport through TNTs is mediated by motor proteins such as kinesin and dynein, a process that requires ATP ^35^. Furthermore, TNT formation itself is driven by actin polymerization, an active and ATP-dependent process ^37^. Consequently, metformin-mediated inhibition of cellular ATP production ^38^ likely impairs both the formation of TNT structures and the active transport of mitochondria across these intercellular connections.

Using a murine model ubiquitously expressing MitoDendra^19^, our study uniquely provides temporal tracking of monocyte recruitment and their fate in the myometrium through adoptive transfer of mitoDendra2-labelled monocytes. The detection of fluorescently labelled mitochondria enabled us to trace monocyte infiltration in the pregnant murine uterus, confirm their differentiation into tissue-resident Macs, characterize their polarization into M1 and M2 subtypes, and to directly demonstrate the mitochondrial transfer from mitoDendra-Macs to mMYOs in pregnant dams at term.

Our findings confirm that monocytes are recruited by the myometrium where they differentiate into Macs within 24 hours. The infiltrated Macs exhibit both M1 and M2 phenotypes; however, close to term a progressive shift towards the M1 phenotype is observed. Importantly, mitochondrial transfer from Macs to MYOs was observed predominantly at full term (GD19) before the onset of labor. This temporality may indicate that TNT-mediated M1-Mac/MYO interactions are established in preparation for labor onset and it is tempting to speculate that they may even provide the essential trigger for myometrial contractions (**Figure 11).**

**Figure 11:**
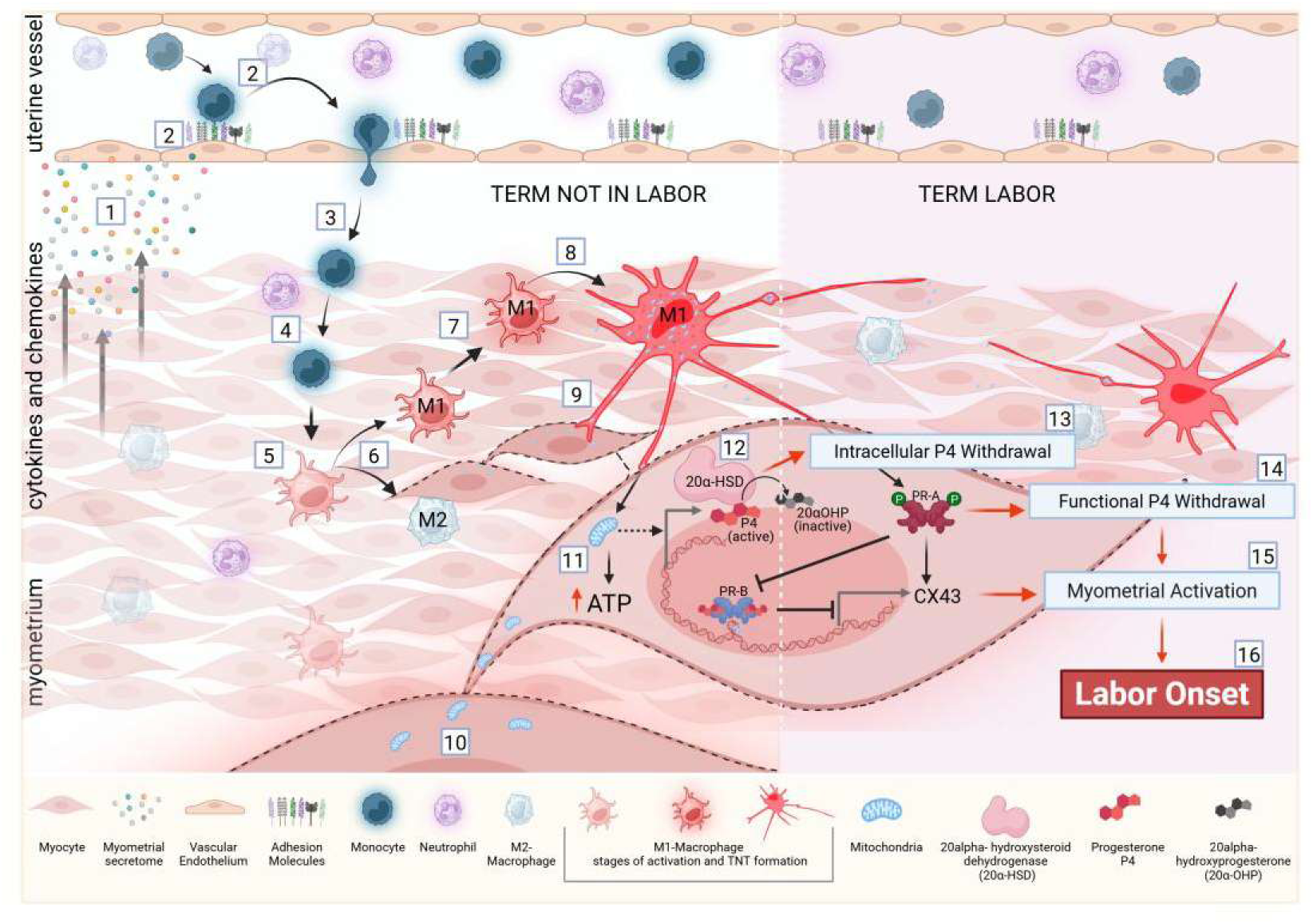
Proposed mechanism of infiltrated monocyte–derived M1 macrophage–driven labor initiation. Schematic illustration depicting the sequence of immunological and metabolic events leading to labor onset. (**1**) Myometrial cells release cytokines and chemokines that (**2**) activate peripheral immune cells and upregulate adhesion molecules on endothelial cells, facilitating (**3**) the recruitment and infiltration of circulating monocytes and neutrophils into the uterus/myometrium. (**4**) Infiltrated monocytes establish residency within the tissue and (**5**) differentiate into macrophages, which subsequently (**6**) polarize into either pro-inflammatory M1 or anti-inflammatory M2 phenotypes. (**7**) At term, M1 macrophages are activated by myocyte-derived signals and (**8**) form tunneling nanotube (TNT) connections with adjacent myocytes. (**9**) Through these TNTs, M1 macrophages transfer mitochondria unidirectionally to myocytes. (**10**) Mitochondria are also shared between myocytes through myocyte-to-myocyte connections. (**11**) Transferred mitochondria enhance ATP production and induce 20α-hydroxysteroid dehydrogenase (20α-HSD) expression, promoting (**12**) the conversion of active progesterone (P4) to its inactive metabolite 20α-hydroxyprogesterone (20α-OHP), resulting in (**13**) intracellular progesterone withdrawal. Concurrently, (**14**) macrophage–myocyte interactions trigger phosphorylation of progesterone receptor PR-A at Ser344/345, stabilizing PR-A, increasing the PR-A:PR-B ratio, and suppressing PR-B–mediated progesterone signaling, thereby contributing to functional progesterone withdrawal. These processes collectively lead to (**15**) myocyte activation, characterized by increased connexin-43 (CX43) expression, culminating in (**16**) the initiation of labor. Illustration created in https://BioRender.com

Collectively, our data suggest that the pregnant uterus actively recruits and functionally repurposes circulating maternal immune cells to facilitate parturition at term. Among these immune cells, M1-Macs appear to play a dominant role in preparing the myometrium for labor onset. They communicate with SMCs, and transfer mitochondria via TNTs, which not only contributes to the induction of P4 withdrawal but also enhances the energetic capacity of the myometrium by increasing ATP availability, thereby supporting the high energy demands of labor. This study fills key gaps in delineating the cellular and molecular interplay between the immune system and the pregnant myometrium and defines the role of M1-Mac-MYO interactions in triggering P4 withdrawal and myometrial activation. We speculate that targeting key processes including Mac infiltration, M1-polarization, TNT formation, M1-Mac-MYO communication and cargo transfer offer novel therapeutic avenues for PTB prevention in high-risk women.

## Acknowledgements

This study was supported by the CIHR Project Grant (PJT-180394) awarded to S.L. and O.S., CIHR (FRN 15608) to A.J. and CIHR (MFE 187836) to A.S. The imaging work was supported by the Lunenfeld-Tanenbaum Research Institute’s Network Biology Collaborative Centre, which is funded by the Canada Foundation for Innovation, the Ontario Government, Genome Canada and Ontario Genomics (OGI-139) and the Nikon Center of Excellence at the Lunenfeld-Tanenbaum Research Institute. L.P. Is a Tier 1 Canada Research Chair in Centrosome Biogenesis and Function. All Illustration in figures are created in https://BioRender.com.

## Author Contributions

L.N., O.S., and S.L. designed the project. L.N., and E.A., performed the research. L.N. and S.S. analyzed the data. A.S. facilitated time-lapse imaging. L.P. provided access to high-content microscopy. A.J. contributed the mouse model. L.N. wrote the manuscript, O.S. and S.L. edited the manuscript.

